# A direct RNA sequencing workflow for simultaneous profiling of tRNA abundance and modifications in mouse liver

**DOI:** 10.64898/2026.10.04.754913

**Authors:** Shenghong Dai, Zhaoyang Tian, Yaxuan Zhang, Xiaojing Li, Xinxin Yi, Ting Qiu, Liu Yang, Jin Ke, Min Chen, Jia Huang, Xinyu Hu, Yanqiang Li, Hu Chen, Tianyuan Zhang

## Abstract

Transfer RNA (tRNA) is highly modified and difficult to profile using reverse-transcription-based sequencing. Here, we established a direct RNA sequencing (tRNA-DRS) workflow to simultaneously characterize tRNA abundance and modification signals in mouse liver. The workflow included small-RNA enrichment, deacylation, splint-adapter ligation, barcoding, reverse transcription, and nanopore sequencing on PromethION with modification-aware basecalling. From 1,678,666 raw reads, 1,421,279 (84.67%) reads were retained after 20–200 nt length filtering, and 17,094 uniquely mapped reads (MAPQ ≥ 10) were obtained. These reads covered 244 tRNA genes and represented all 23 tRNA isotypes. tRNA abundance was strongly skewed, with Asp, Gly, Ser, and Lys accounting for 56.6% of uniquely mapped reads. Read-body coverage indicated that most uniquely mapped reads spanned substantial portions of tRNA molecules (mean 71.1%, median 75.9%; 83.3% reads covering ≥50% of the reference). Modification analysis identified 41,013 candidate sites, of which 711 passed stringent filtering (modification fraction ≥50%, coverage ≥5), spanning eight modification types and dominated by pseudouridine and m5C. Filtered sites showed position-specific clustering and isotype-dependent patterns. Together, these results support tRNA-DRS as a practical single-assay framework for simultaneous tRNA expression and epitranscriptomic profiling in mammalian tissue.

## Introduction

Transfer RNA sits at an odd position in the RNA world: short, abundant, and easy to overlook, yet essential for protein synthesis. Each tRNA carries an anticodon and the corresponding amino acid, and the cell depends on this match to translate the genetic code with acceptable accuracy. tRNA does not simply ferry amino acids, though. Recent work connects tRNA abundance, codon choice, and processing into tRNA-derived fragments to cell growth, metabolism, and disease ^1–3^. The list of disease associations keeps growing, from cancer to mitochondrial and neurological disorders ^1,4^.

Part of the difficulty in studying tRNA lies in its modification density. A single tRNA molecule can carry a dozen or more chemical modifications, and more than one hundred modification types have been found across the tRNA pool ^1,2^. These marks tune codon-anticodon pairing, folding, stability, and aminoacylation; their loss can change translation without changing the sequence ^2,4^. Some tRNA-modifying enzymes are now considered candidate drug targets ^4^.

Standard sequencing approaches do not see these marks directly. Most RNA-seq pipelines start with adapter ligation, reverse transcription, and amplification, and reverse transcriptase tends to pause or misread at modified bases. Mass spectrometry can quantify modified nucleosides but needs microgram-scale RNA and loses sequence context. Antibody- and chemical-based methods only cover certain modifications. None of them gives a full, sequence-resolved picture of a tRNA molecule as it exists in the cell.

Nanopore direct RNA sequencing takes a different route. Native RNA is threaded through a protein pore, and the ionic current produced by each nucleotide depends on its chemical identity, so modified bases leave a readable trace in the raw signal ^5,6^. Because the RNA is not amplified, the modification state is preserved. Nanopore direct RNA-seq was used to mapping internal m6A, m5C ^7,8^, Nm ^9,10^ and revealed their functions in mRNA, such as stabilizing mRNA ^11^. Recent reviews cover the technology and its growing use for transcriptome- and epitranscriptome-scale studies ^7,8^. Systematic work with synthetic standards has also clarified which modifications can be detected and how reliably ^12^.

tRNA used to be a hard target for nanopore sequencing because the molecule is only about 76 nt long and folds into a stable cloverleaf, which makes end-to-end adapter ligation difficult. A splint-ligation protocol changed that: adapters are annealed to both ends of deacylated tRNA, and the resulting library can be sequenced directly on a nanopore flowcell ^13^. The approach has been used to quantify tRNA abundance and modifications, applied to cancer cell models ^14^, and extended by dedicated modification callers such as QutRNA2 ^15^.

Here, we tested a tRNA direct RNA sequencing workflow using one mouse liver sample. The pipeline covered tRNA enrichment, deacylation, splint adapter ligation, barcoding, reverse transcription, and nanopore sequencing. This study is not intended to address liver biology; it focuses on whether the method can generate usable tRNA reads for downstream abundance and modification analysis.

## 2. Methods

### 2.1 Experimental procedures

#### 2.1.1 Total RNA extraction and tRNA enrichment

Total RNA was extracted from mouse liver with TRIzol ^16^. 10 µg per sample was used for tRNA enrichment. The small RNA fraction containing tRNA was enriched with the Monarch Spin RNA Cleanup Kit (New England Biolabs, T2040), following the large/small RNA separation protocol. The sample was mixed with 1 volume of Buffer BX and 1/2 volume of ethanol, then loaded onto a column. Large RNA (>200 nt) was retained on the column, whereas small RNA remained in the flow-through. The flow-through was re-loaded onto a fresh column, washed, and eluted, yielding the enriched small RNA fraction (<200 nt, containing tRNA) ^17^.

#### 2.1.2 tRNA library construction and Nanopore sequencing

The enriched small RNA was deacylated in 90 µL of deacylation buffer (final volume 100 µL) at 37°C for 30 min ^18^, purified with the RNA Clean & Concentrator-5 kit (Zymo Research, R1015/R1016), and eluted in 8 µL of nuclease-free water. At least 250 ng of deacylated tRNA per sample, quantified with the Qubit HS RNA Assay Kit (Thermo Fisher), was used for library construction. Library construction followed the splint-ligation strategy established for nanopore tRNA sequencing ^13,19^.

Splint adapters A and B were annealed in 10 µL reactions containing 1 µL each of annealing buffers AB1 and AB2, 3.75 µL each of the adapters, and 0.5 µL of murine RNase inhibitor (New England Biolabs, M0314L). Annealing was performed by heating to 75°C for 15 s and cooling to 25°C at 0.1°C/s. The annealed splint adapter was ligated to 250 or 500 ng of deacylated tRNA in 25 µL reactions containing 10 µL of PEG 8000, 2.5 µL of ligation buffer (T1), 2 µL of ligase (T1), and 0.5 µL of RNase inhibitor, at 4°C overnight. Ligation products were purified with Agencourt AMPure XP beads (Beckman Coulter, A63881), washed twice with freshly prepared 70% ethanol, and eluted in 9 µL of nuclease-free water. The splint-ligation strategy overcomes the difficulty of end-to-end adapter ligation on short, stably folded tRNA molecules.

Each sample was then ligated to a unique pre-annealed barcoded adapter (BC1–BC6; annealing as for the splint adapters) in 15 µL reactions at room temperature for 10 min. Reverse transcription was performed in 24.5 µL master mixes containing 14.5 µL of nuclease-free water, 8 µL of reverse-transcription buffer T3, and 2 µL of dNTPs, with 0.5 µL of reverse transcriptase (T3), at 60°C for 30 min followed by 85°C for 5 min. The cDNA was purified with AMPure XP beads and eluted in 8.5 µL of nuclease-free water.

Barcoded samples were pooled to 20 ng of total nucleic acid in 23 µL (equal amounts of up to six samples, quantified with the Qubit RNA Assay Kit, Thermo Fisher). The RNA ligation adapter was ligated in a 40 µL reaction containing NEBNext Quick Ligation Reaction Buffer, 6 µL of RNA Ligation Adapter (SQK-RNA004), and 3 µL of T4 DNA Ligase, at room temperature for 10 min. The library was purified with Agencourt RNA Clean XP beads (Beckman Coulter, A63987), washed, and eluted in 13 µL of RNA Elution Buffer. A 100 µL volume of SB (SQK-RNA004) and 68 µL of LIS were added to the eluate. The library was loaded immediately onto an FLO-PRO004RA flow cell and sequenced on a PromethION instrument using the Direct RNA Sequencing Kit (SQK-RNA004, Oxford Nanopore Technologies). Sequencing was performed by Wuhan Benagen Technology Co., Ltd.

### 2.2 Computational analysis of tRNA direct RNA sequencing

#### 2.2.1 Basecalling

Raw signals were basecalled with dorado (v0.9+), using the RNA004 130-bp/s model sup@v5.2.0 with four modification bundles (m5C_2OmeC, inosine_m6A_2OmeA, pseU_2OmeU, 2OmeG), which decode eight tRNA modifications: m5C, Cm, Gm, pseudouridine (Ψ), Um, m6A, Am, and inosine. Basecalling removed adapters and poly(A) tails. Modification information was written to the output BAM as MM/ML (and MN) tags for downstream alignment and quantification. The performance of these models for tRNA modification detection has been evaluated previously ^20,21^.

#### 2.2.2 Read filtering

Full-length tRNAs are much shorter than most cellular RNAs, whereas DRS of native RNA is contaminated by rRNA, mRNA, and non-tRNA fragments. To enrich for tRNA and reduce alignment noise, we retained reads of 20–200 nt (preserving MM/ML tags) and discarded unmapped, secondary, and supplementary alignments.

#### 2.2.3 Alignment

The alignment reference comprised mature tRNA gene sequences from the GtRNAdb annotation of the mouse genome (Mus musculus, GRCm38/mm10) ^17^, covering all amino-acid families (isotypes). A high-confidence subset of the same annotation was used for robustness assessment. Alignments were generated with bwa mem using the -x ont2d preset (-C -W 13 -k 6 -T 20), which applies low mismatch and indel penalties and improves the alignment rate of low-abundance tRNAs. The -C option carries the MM/ML/MN tags into the alignment, preserving modification information.

#### 2.2.4 Quantification

tRNA expression counts per gene were generated from primary alignments with a custom quantification script. Counts of uniquely mapped reads (MAPQ ≥ 10) formed the main quantification set, together with all primary-alignment reads. Counts were aggregated by amino-acid family into isotype-level counts and by amino-acid–anticodon combination into anticodon-level counts. Per-gene annotation (GtRNAdb ID, confidence, amino-acid family, anticodon, genomic coordinates, isotype and structure scores) was output with the expression counts.

#### 2.2.5 Modification detection

Modification detection was performed with modkit on the MM/ML tags in the post-alignment BAM. Reads were filtered to 20–200 nt (retaining MM/ML tags), converted to FASTQ with samtools fastq -T MM,ML,MN, realigned with bwa mem as described above, and filtered to MAPQ ≥ 10. modkit pileup was then run to compute per-position modification fractions. modkit is an open-source tool for modification calling and summarization from nanopore data.

At each reference position, the modification fraction was the proportion of reads supporting the modification among all effective reads at that position. A site was called a high-confidence modification when the fraction was ≥ 50% and the coverage was ≥ 5. All sites were summarized with coverage and modification fraction; high-confidence sites were reported separately and grouped by modification type.

#### 2.2.6 Full-length coverage and positional profiling

For each uniquely mapped read, reference coverage was the fraction of the reference length spanned by the read, aggregated per gene and per isotype to obtain mean and median coverage and the fraction of reads with ≥90%, ≥80%, or ≥50% coverage. These metrics were used to assess full-length tRNA read quality.

The modification landscape was summarized at two levels. First, the 5′→3′ positional profile: high-confidence modifications of each gene were aggregated by tRNA position to compute the modification rate per position. Second, the isotype × modification-type matrix: the number of high-confidence sites within each amino-acid-family × modification-type combination was counted to reveal modification preferences across tRNA families.

## Results

### 3.1 The tRNA-DRS workflow applied to the size-selected small-RNA fraction of mouse liver

To profile tRNA expression and modifications in mouse liver, we applied the tRNA-DRS workflow to the size-selected small-RNA fraction (<200 nt) from a single library, with reads sequenced on a PromethION instrument and basecalled with modification models. Nanopore direct RNA sequencing (DRS) enables direct, amplification-free analysis of native tRNA molecules, including their covalent modifications ^13,19,22^. The run generated 1,678,666 raw reads. After length filtering to 20–200 nt, 1,421,279 reads (84.67%) were retained (Fig. 1A), with read lengths consistent with the expected small-RNA size profile (Fig. 1B). Following alignment to the GtRNAdb-annotated mouse tRNA reference (GRCm38/mm10), 17,094 reads (1.02%) mapped uniquely to tRNA genes with MAPQ ≥ 10 (Fig. 1D). These results establish the tRNA-DRS workflow for mouse liver and quantify the low abundance of tRNAs within the small-RNA transcriptome; because the present dataset derives from a single library, the quantitative findings below should be regarded as a pilot assessment.

**Fig. 1.**
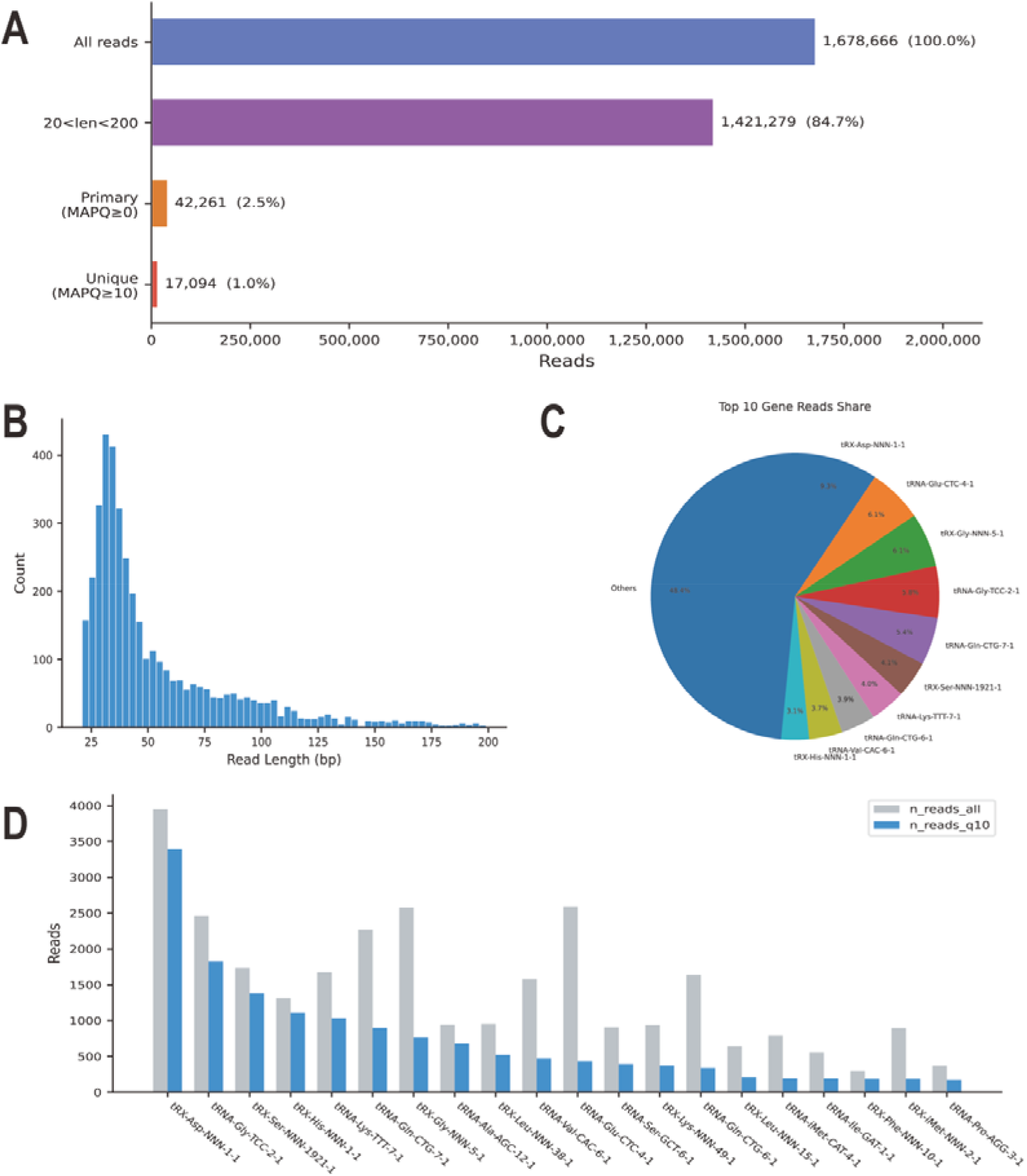
Library composition and gene-level coverage. A, Bar chart of reads retained at each filtering step. B, Histogram of read lengths for the length-filtered reads (20–200 nt; n = 1,421,279). C, Pie chart of the read share of the ten most abundant tRNA genes (n = 42,261 primary alignments; MAPQ ≥ 0). D, Grouped bar chart of total (gray) versus unique (blue) alignments per gene for the 20 most abundant genes (n = 731 annotated tRNA genes; primary, MAPQ ≥ 0; unique, MAPQ ≥ 10).

### 3.2 All tRNA isotypes are detected despite a low unique-mapping fraction

Despite the low yield of uniquely mapped reads, tRNA-DRS recovered a broad spectrum of tRNA genes. At any mapping quality, 467 of 731 annotated mouse tRNA genes (63.9%) were covered by at least one read, and 244 genes (33.4%) were covered by uniquely mapped reads (MAPQ ≥ 10;

Fig. 1D). All 23 isotypes (amino-acid families) were represented among the covered genes, and coverage depth was not merely token: 17 of 23 isotypes were covered by at least 50 uniquely mapped reads (median 297 reads per isotype, range 0–3,400; Fig. 1D). Gene-level coverage varied widely, ranging from single reads to hundreds of uniquely mapped reads per gene, consistent with large differences in tRNA gene copy number and expression level across the genome ^13^.

### 3.3 tRNA expression is strongly skewed toward a small subset of isotypes and anticodons

tRNA expression was strongly skewed at the isotype level: four isotypes—Asp, Gly, Ser, and Lys—accounted for 56.6% of all uniquely mapped reads, led by Asp (3,400 reads) and Gly (2,735 reads) (Fig. 2A). At the gene level, uniquely mapped reads were similarly concentrated: the ten most abundant tRNA genes accounted for 70.6% of all uniquely mapped reads (Fig. 1C). This skew was retained after normalization: the same four isotypes showed the highest expression in reads per million (RPM, 198,900–110,741) and represented ~41% of total RPM. At the anticodon level, Asp-ATC (3,398 reads), Gly-TCC (1,842), Lys-TTT (1,423), Ser-CGA (1,380), and Gln-CTG (1,235) were the most abundant, together representing 54.3% of uniquely mapped reads (Fig. 2B). This skewing toward a small number of isotypes and anticodons is qualitatively consistent with the dominance of high-copy tRNA genes in mammalian tRNA transcriptomes ^13,14^. tRNA species dominated the decoding capacity of mouse liver.

**Fig. 2.**
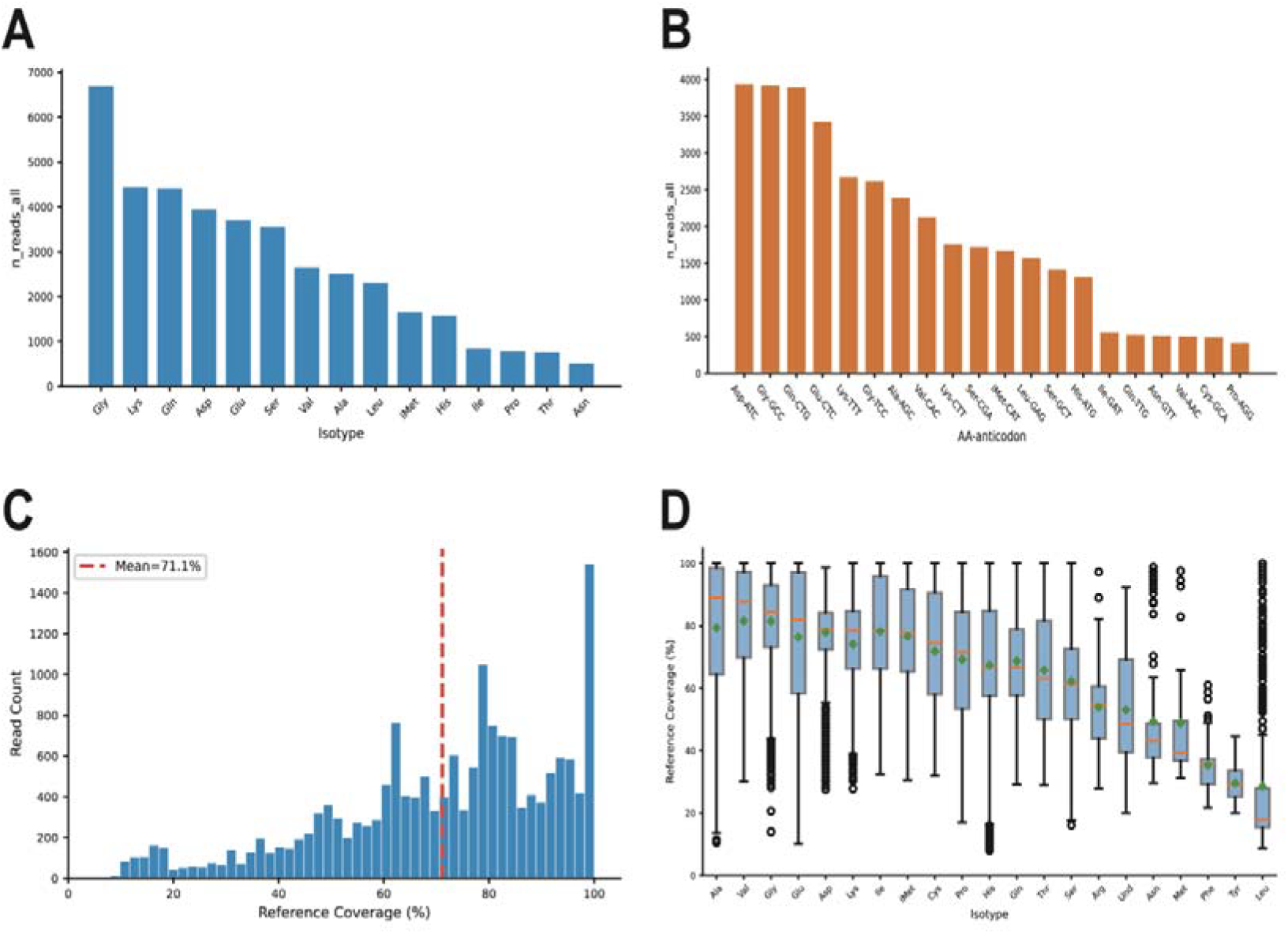
Isotype- and anticodon-level expression and full-length coverage. A, Bar chart of uniquely mapped reads (MAPQ ≥ 10) per isotype (n = 23 isotypes). B, Bar chart of uniquely mapped reads per anticodon. C, Histogram of per-read reference-body coverage, i.e. the fraction of the tRNA reference spanned by each read (n = 17,094 uniquely mapped reads); vertical lines indicate the mean and median. D, Box plots of per-read reference coverage by isotype.

### 3.4 Most uniquely mapped reads span the majority of the tRNA body

To determine whether uniquely mapped reads represented full-length tRNAs, we computed for each read the fraction of the reference tRNA length covered. Mean and median reference coverage were 71.1% and 75.9%, respectively (Fig. 2C). The large majority of reads (83.3%; 14,233 reads) covered at least 50% of the tRNA body, whereas 40.6% (6,933 reads) and 22.6% (3,862 reads) covered at least 80% and 90%, respectively (Fig. 2C). Coverage distributions were broadly similar across isotypes (Fig. 2D). These results indicate that most uniquely mapped reads encompass the majority of the tRNA sequence, providing a suitable substrate for full-length modification profiling ^15^, although a tail of short reads may reflect partially degraded or truncated tRNA species.

### 3.5 tRNA-DRS detects stringently filtered modification sites across multiple modification types

Using four modification model bundles, we identified 41,013 candidate modification sites across the tRNA reference. Application of a pre-defined filter (modification fraction ≥ 50% and coverage ≥ 5) retained 711 sites (Fig. 3A). The filtered sites spanned all eight modification types, with pseudouridine (Ψ; 195 sites) and 5-methylcytosine (m5C; 176 sites) being the most abundant and together accounting for 52.2% of all sites, followed by 2′-O-methylated nucleosides (Um, 102; Gm, 87; Cm, 32; Am, 30), N6-methyladenosine (m6A; 65), and inosine (24) (Fig. 3A). The number of filtered sites per gene varied widely, with a subset of genes carrying more than ten sites (Fig. 3B). This spectrum is broadly consistent with the known tRNA modification repertoire in bacteria and yeast ^21,23–25^ and with the global impact of pseudouridylation on other modification types reported in nanopore DRS studies ^26^.

**Fig. 3.**
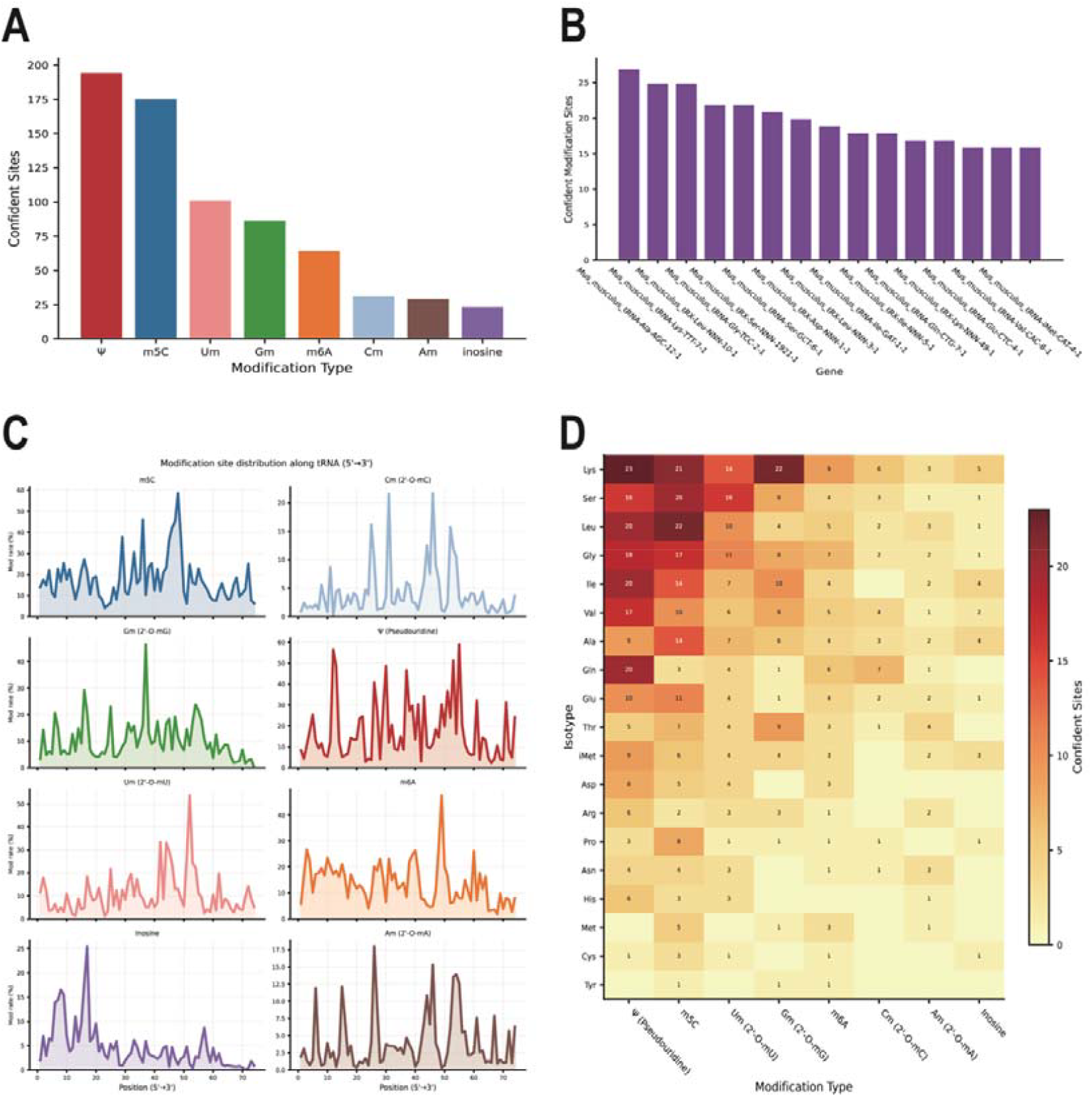
Modification sites across types, genes, positions and isotypes. A, Bar chart of stringently filtered modification sites (modification fraction ≥ 50% and coverage ≥ 5; Methods 2.2.4) by modification type. B, Distribution of the number of stringently filtered sites per gene (n = 244 uniquely mapped genes). C, Number of stringently filtered sites at each reference position (positions 1–74). D, Heatmap of stringently filtered sites per isotype and modification type (rows, isotypes; columns, modification types).

### 3.6 Filtered modification sites cluster at discrete positions and differ across isotypes

Among the stringently filtered sites, the modification landscape appeared non-uniform along the tRNA sequence. Positional profiling across tRNA positions 1–74 showed that filtered sites were concentrated at discrete positions (Fig. 3C), qualitatively consistent with the known clustering of modified nucleosides in the anticodon loop and TΨC arm ^21,27,28^, although we did not test this against a random-position null model. In addition, the isotype × modification-type matrix revealed distinct modification preferences among amino-acid families: individual isotypes differed in both the number and the combination of modification types detected (Fig. 3D). These patterns are consistent with type-specific regulation of tRNA stability and decoding reported in prior work ^24,26^ and, together with the expression data, illustrate the potential of tRNA-DRS as a single-assay approach for simultaneous expression and epitranscriptomic profiling of tRNA families.

## Acknowledgements

The experimental design and operation methods of this study have been reviewed and approved by the Experimental Animal Ethics Committee of Huanghuai University (Approval No.: HHU202603090028).

